# Sleep and circadian rhythm disruption by NPTX2 loss of function

**DOI:** 10.1101/2023.09.26.559408

**Authors:** Seung-Eon Roh, Meifang Xiao, Ana Delgado, Chuljung Kwak, Alena Savonenko, Arnold Bakker, Hyung-Bae Kwon, Paul Worley

## Abstract

Sleep and circadian rhythm disruption (SCRD) is commonly observed in aging, especially in individuals who experience progressive cognitive decline to mild cognitive impairment (MCI) and Alzheimer’s disease (AD). However, precise molecular mechanisms underlying the association between SCRD and aging are not fully understood. Orexin A is a well-characterized “sleep neuropeptide” that is expressed in hypothalamic neurons and evokes wake behavior. The importance of Orexin is exemplified in narcolepsy where it is profoundly down-regulated. Interestingly, the synaptic immediate early gene NPTX2 is co-expressed in Orexin neurons and is similarly reduced in narcolepsy. NPTX2 is also down-regulated in CSF of some cognitively normal older individuals and predicts the time of transition from normal cognition to MCI. The association between Orexin and NPTX2 is further evinced here where we observe that Orexin A and NPTX2 are highly correlated in CSF of cognitively normal aged individuals and raises the question of whether SCRD that are typically attributed to Orexin A loss of function may be modified by concomitant NPTX2 down-regulation. Is NPTX2 an effector of sleep or simply a reporter of orexin-dependent SCRD? To address this question, we examined NPTX2 KO mice and found they retain Orexin expression in the brain and so provide an opportunity to examine the specific contribution of NPTX2 to SCRD. Our results reveal that NPTX2 KO mice exhibit a disrupted circadian onset time, coupled with increased activity during the sleep phase, suggesting difficulties in maintaining states. Sleep EEG indicates distinct temporal allocation shifts across vigilance states, characterized by reduced wake and increased NREM time. Evident sleep fragmentation manifests through alterations of event occurrences during Wake and NREM, notably during light transition periods, in conjunction with an increased frequency of sleep transitions in NPTX2 KO mice, particularly between Wake and NREM. EEG spectral analysis indicated significant shifts in power across various frequency bands in the wake, NREM, and REM states, suggestive of disrupted neuronal synchronicity. An intriguing observation is the diminished occurrence of sleep spindles, one of the earliest measures of human sleep disruption, in NPTX2 KO mice. These findings highlight the effector role of NPTX2 loss of function as an instigator of SCRD and a potential mediator of sleep disruption in aging.

## Introduction

Sleep and circadian rhythm disruption (SCRD) is a prominent characteristic of aging and neurological diseases, including Alzheimer’s disease (AD) and mild cognitive impairment (MCI) (Musiek, Xiong, and Holtzman 2015; Guarnieri et al. 2012). Sleep disturbances, such as sleep fragmentation, daytime sleepiness, and circadian disruption, are commonly observed in individuals with AD and MCI (Lim et al. 2013; Kaneshwaran et al. 2019; Lee et al. 2007; Spira et al. 2018; Pak et al. 2020). Moreover, SCRD often occurs before cognitive decline (Spira et al. 2018) and is believed to worsen disease progression (Lloret et al. 2020). Importantly, SCRD has been recognized as a potential modifiable risk factor for AD (Minakawa, Wada, and Nagai 2019). Despite the importance of earlier and more accurate detection and intervention of the diseases, the specific molecular mechanisms underlying SCRD in aging and the development of neurological diseases are not yet fully understood.

NPTX2 is a synaptic protein and an immediate early gene predominantly expressed in pyramidal neurons (Xu et al. 2003). In response to specific patterns of stimuli, NPTX2 undergoes exocytosis and specifically accumulates at excitatory synapses onto parvalbumin interneurons (PV-IN) (Chang et al. 2010; Xiao et al. 2021). This process enhances the excitatory drive of inhibition by facilitating the clustering of AMPA receptors, thereby playing a crucial role in regulating the balance between excitation and inhibition (Chang et al. 2010; Pelkey et al. 2015). Notably, NPTX2 levels are diminished in AD brain and in CSF of individuals with AD and MCI (Xiao et al. 2017). Furthermore, the concentration of NPTX2 in the CSF is associated with cognitive performance and hippocampal volume in AD and MCI patients, making it an effective biomarker for these conditions (Xiao et al. 2017; Galasko et al. 2019). Notably, NPTX2 reduction in the CSF can even predict the progression from normal cognition to MCI, suggesting that NPTX2 is one of the earliest measures useful for the early detection of diseases (Soldan et al. 2023). Additionally, reduced levels of NPTX2 in the CSF have been observed in individuals with recent-onset schizophrenia, and its concentration correlates with cognitive performance (Xiao et al. 2021). Biochemical analysis of schizophrenia and bipolar disorder patient tissues also revealed reduced levels of NPTX2 in synaptosome fraction (Aryal et al. 2023). Importantly, sleep is disrupted to various degrees in each of these conditions.

Several studies have provided evidence supporting the regulation of NPTX2 expression during sleep-wake cycles. It was shown that mRNA levels of NPTX2 in the neocortex and cerebellar cortex exhibit an upregulation during wakefulness (Cirelli, Gutierrez, and Tononi 2004). Also, sleep deprivation can induce NPTX2 expression, along with other immediate early genes such as Arc, Homer1a, and BDNF (Thompson et al. 2010). Interestingly, NPTX2 expression is induced in the cells, where Homer1a is induced by sleep deprivation (Maret et al. 2007). We have demonstrated that the exocytosis and shedding of synaptic NPTX2 are regulated over the sleep-wake cycle, and sleep deprivation enhances the synaptic levels of NPTX2 in mice and the CSF levels of NPTX2 in healthy individuals (Xiao et al. 2021). These observations suggest that extended periods of wakefulness promote both the exocytosis and shedding of NPTX2 and indicate that NPTX2 is influenced by sleep-wake cycles. Accordingly, NPTX2 may play a role in the dynamic processes associated with sleep and wakefulness.

Orexin, also known as hypocretin, is a prominent wake-promoting neuropeptide expressed in lateral hypothalamic neurons. The Orexin knockout and Orexin degeneration models display difficulties sustaining wakefulness and evidence of sleep fragmentation albeit relatively normal total amounts of wakefulness (Diniz Behn et al. 2010; Kantor et al. 2009). Autoimmune loss of Orexin results in narcolepsy, which is characterized by excessive daytime sleepiness, cataplexy (sudden loss of muscle tone), and disrupted REM sleep (Dauvilliers, Arnulf, and Mignot 2007). Interestingly, NPTX2 is highly enriched in the Orexin neurons (Dalal et al. 2013; Reti et al. 2002) and human studies have revealed that NPTX2, along with dynorphin and Orexin, are concurrently lost in the Orexin neurons of the lateral hypothalamus in individuals with narcolepsy (Crocker et al. 2005). Additionally, NPTX2 has been suggested to play a significant role in synaptic structural plasticity within Orexin neurons during the sleep-wake cycle (Appelbaum et al. 2010). Overexpression of NPTX2 in Orexin neurons abolished this synaptic plasticity, leading to melatonin resistance. These findings suggest a potential interaction between Orexin and NPTX2 in regulating sleep.

Our study here was prompted by the previously unreported positive correlation of NPTX2 and Orexin A in the CSF of normal, aged individuals (Figure 1A). Thus, individuals with low levels of Orexin A also have low levels of NPTX2. This observation, together with prior basic and translational studies, raises the question of whether SCRD that may be presumed to occur as a result of reduced Orexin might be modified by reduced NPTX2. To address the individual contribution of NPTX2 loss of function in SCRD, we first confirmed that Orexin A is not reduced in NPTX2 KO mice(Figure 1B) and then examined sleep and circadian rhythm by employing wheel-running behavior assessment, sleep EEG, and EEG analysis in NPTX2 KO mice. Studies demonstrate that NPTX2 is a critical effector of sleep and its loss of function contributes to specific features of SCRD.

**Figure 1.**
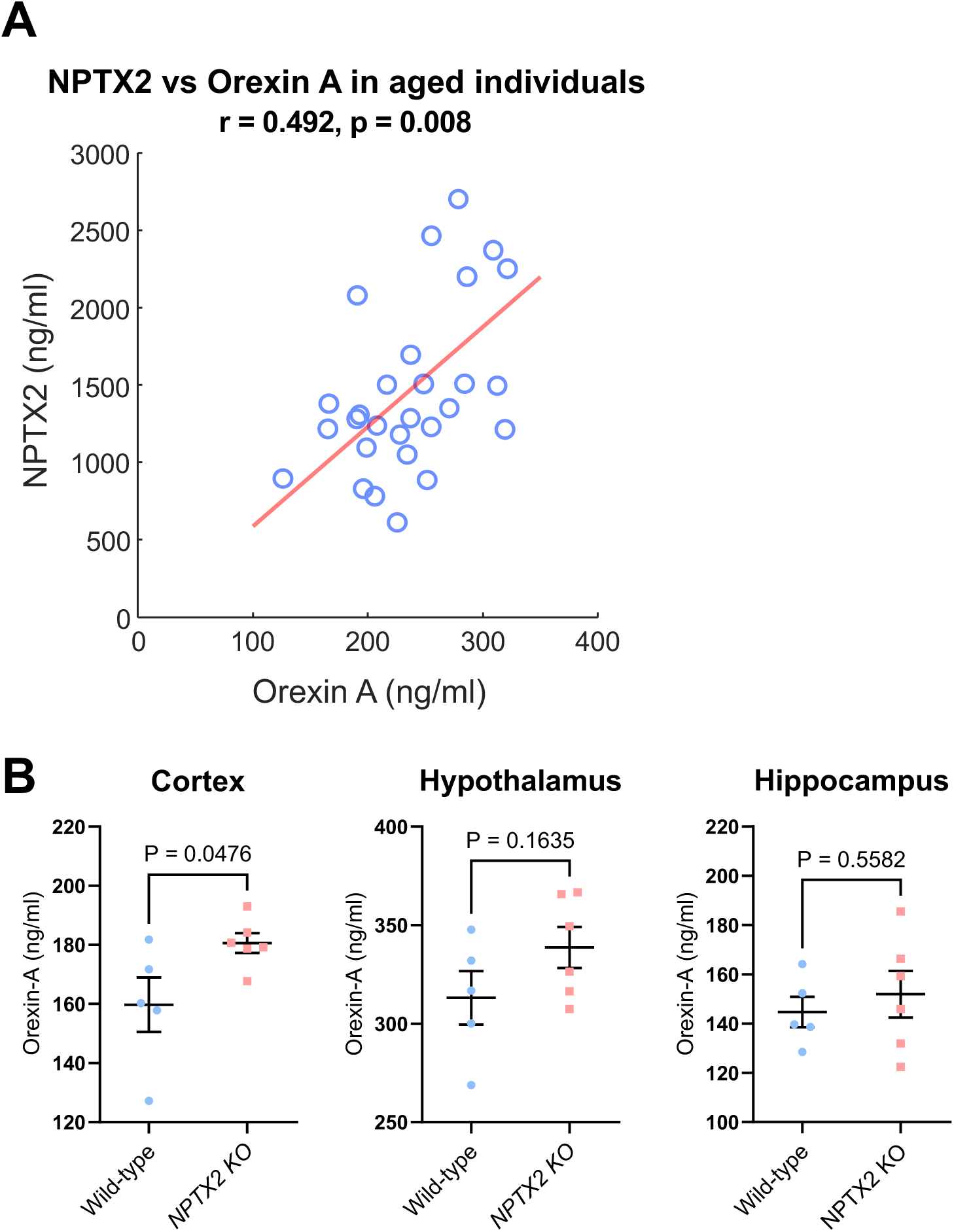
Correlation of NPTX2 and Orexin in the CSF of normal aging and Orexin levels in NPTX2 KO mice. **(A)** Spearman regression analysis between the level of CSF NPTX2 and Orexin A in normal individuals. n = 29 human individuals. r and p values were indicated. **(B)** The concentration of Orexin in lysates of the cortex, hypothalamus and hippocampus between wild-type and NPTX2 KO mice, determined by Orexin A ELISA. N = 5 wild-type and 6 NPTX2 KO. An unpaired t-test was performed. P values were indicated.

## Results

### Correlated levels of NPTX2 and Orexin A in the CSF of normal-aged individuals

NPTX2 is enriched in Orexin neurons in the lateral hypothalamus where NPTX2 is lost alongside the Orexin in the narcolepsy brain (Crocker et al. 2005; Reti et al. 2002). Together with these, the observation that abolished rhythmicity of NPTX2 expression impairs the structural plasticity of Orexin neuronal axon (Appelbaum et al. 2010) suggests associated functionality by NPTX2 and orexin. Using ELISA, we measured NPTX2 and Orexin in the CSF of 29 normal individuals with an average age of 69.2 (Alm et al. 2022) and found a strong positive correlation (Figure 1A, r = 0.492, p = 0.008). This suggests that the Orexin circuit may depend on the NPTX2 function in aging. To determine the effect of NPTX2 loss of function in Orexin level in a mouse model, we performed ELISA with brain lysates of NPTX2 KO mice – cortex, hippocampus, and hypothalamus. Interestingly, however, we found no reduction in Orexin contents in any part of the NPTX2 KO mice brain that were tested (Figure 1B). Rather, there was a significant increase in Orexin levels in the cortex of NPTX2 KO mice (Figure 1B, P = 0.0476). These suggest a correlated but complicated co-functionality between NPTX2 and Orexin in normative aging.

### Unstable circadian onset time and activity in NPTX2 KO mice

With an expectation that NPTX2 KO mice will have similar phenotypes with Orexin KO mice, we set out to characterize sleep and circadian rhythm phenotypes in NPTX2 KO mice. As Orexin KO mice have difficulty maintaining wakefulness (Diniz Behn et al. 2010), we first asked whether NPTX2 KO mice have disrupted circadian rhythm and activity patterns between light and dark phases, using a wheel-running behavior assessment. This analysis involved monitoring the activity of 10 wild-type and 10 NPTX2 KO mice during the 2 weeks of normal light-dark cycle, 2 weeks of complete darkness, and another 2 weeks of light-dark cycle over 6 weeks. In wild-type mice, we observed a clear distinction in activity patterns between the light-on and light-off phases, with the onset time consistently shifting to the left, indicating a normal circadian period slightly shorter than 24 hours (Figure 2A, B, D). The average period of NPTX2 KO mice under complete darkness was comparable with that of wild-type mice (Figure 2B), suggesting that circadian rhythm may be functional in NPTX2 KO mice. Also, like in wild-type mice, NPTX2 KO mice were re-entrained to the light-dark schedule after the complete darkness schedule, indicating that they respond to light. However, NPTX2 KO mice exhibited frequent activity during the light phase under the light-dark cycle, and remarkably, the onset time appeared irregular over the course of days (complete darkness, Figure 2A, D, and E). Furthermore, despite individual variations (Supplemental Figure 1), there was a significant increase in the covariance of onset time over days in NPTX2 KO mice (Figure 2F, P < 0.05). These findings strongly suggest that NPTX2 KO mice face challenges transitioning between day and night, causing unstable timing of circadian onset.

**Figure 2.**
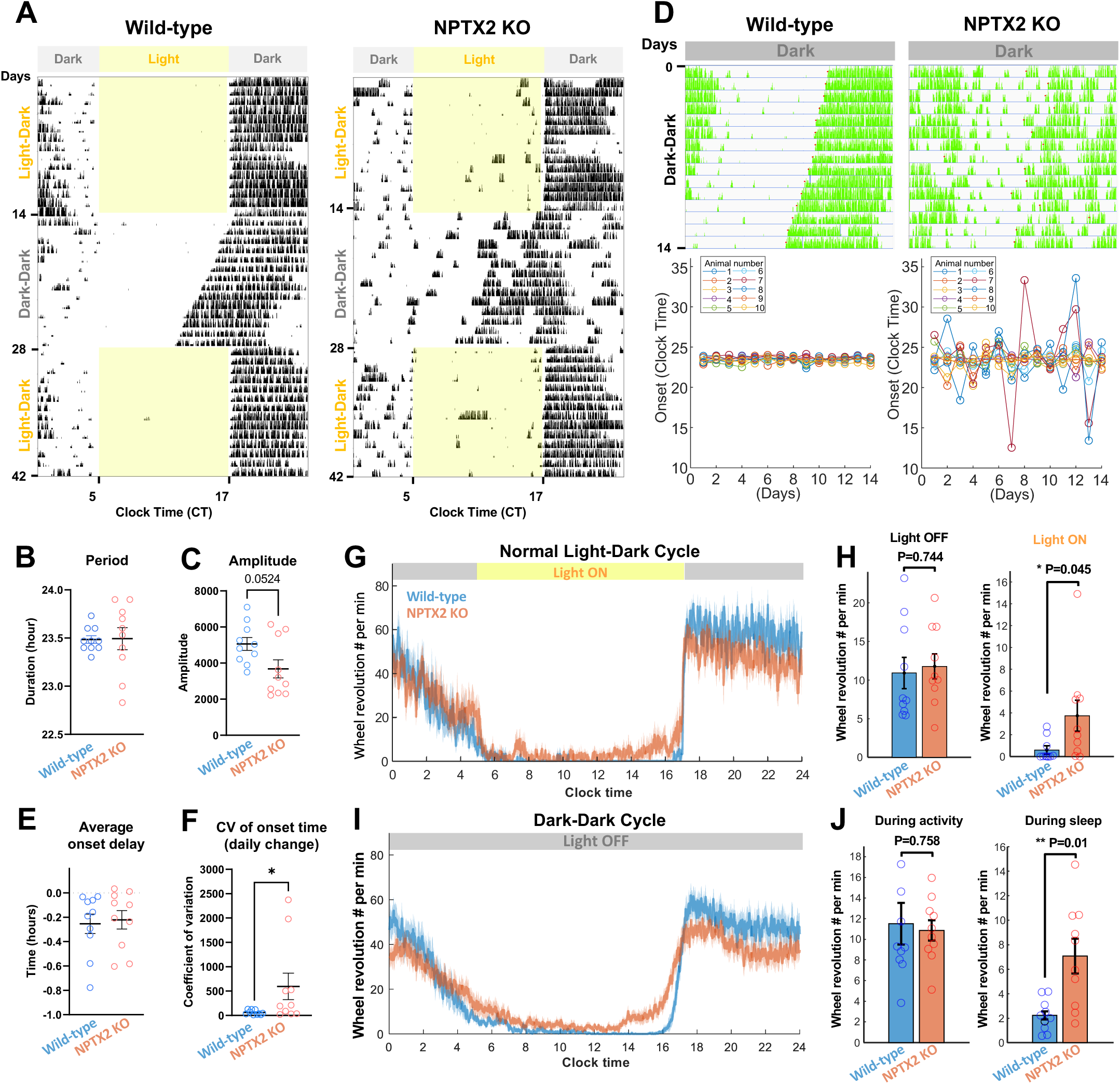
Disrupted circadian rhythm, onset time and activity profile in NPTX2 KO mice. **(A)** Representative actogram of the wheel-running activity of wild-type and NPTX2 KO mice. Each row represents activity level per minute during a day across the indicated light-dark schedule. During 1-14 and 29-42 days, a normal LD cycle was applied (light on from 5 to 17 CT), while during 15-28 days, light was completely off. **(B-C)** Comparison of the period and amplitude (± SEM) between wild-type and NPTX2 KO mice. Mann-Whitney U test was performed to compare Wild-type and NPTX2 KO mice. **(D)** Representative actogram with onset detection (red) in wild-type and NPTX2 KO mice during DD schedule. Each row represents the activity level per minute each day (upper). The circadian onset time over 14 days was presented in graphs (lower). **(E-F)** Averaged onset delay and covariance (CV) of onset time across 14 days of DD period in wild-type and NPTX2 KO mice. **(G)** Averaged number of wheel revolutions (± SEM) across 14 days of LD schedule. **(H)** Comparison of wheel revolution number between wild-type and NPTX2 KO mice in light off and light on phase. Unpaired t-test was performed to compare wild-type and NPTX2 KO mice. P values were indicated. **(I)** Averaged number of wheel revolutions (± SEM) across 14 days of DD schedule. **(J)** Comparison of wheel revolution number between wild-type and NPTX2 KO mice during activity and during sleep. Unparied t-test was performed to compare between wild-type and NPTX2 KO mice. P values were indicated. N = 10 wild-type and 10 NPT2 KO animals.

Consistent with the reduced amplitude of activity observed in NPTX2 KO compared to wild-type mice (Figure 2C, P = 0.0524), further analysis of the activity levels under the light-dark cycle revealed a significant increase in wheel-running activity during the light phase in NPTX2 KO mice (Figure 2 G-J, P < 0.05). This effect was more pronounced during the sleep phase under the complete darkness schedule (Figure 2 I-J). These results indicate that NPTX2 KO mice struggle to maintain sleep. Taken together, these results indicate that the circadian rhythm regulation of NPTX2 KO mice is relatively normal, while frequent activity during the sleep phase and irregular circadian onset timing suggests unstable maintenance of sleep and/or wakefulness, requiring more intricate analysis of different vigilance states.

### Altered time of vigilance states, and sleep-wake fragmentation in NPTX2 KO mice

For a deeper analysis of sleep-wake states in NPTX2 KO mice, we have performed EEG recording using the Neurologger, a wireless electrophysiological data logger. The assessment in 11 wild-type and 12 NPTX2 KO mice involved sleep architecture analysis among the three different vigilance states: wake, NREM, and REM, over a 24-hour period (12 hours of light on and 12 hours of light off) by implementing semi-automatic sleep annotation using custom-written python script. Our analysis in 1-hour bins revealed significant differences in the time spent in these states between the wild-type and NPTX2 KO mice in some time ranges (Figure 3 C). Specifically, we observed a reduction in wake time and an increase in NREM sleep time, especially near the transition periods (Figure 3C, P < 0.05). When quantifying the vigilance states over the 12 hours of light on, 12 hours of light off, and the total 24-hour period, we found a modest reduction in wake time (light-off: 53.75 ± 3.05 compared to 61.27 ± 2.52; total: 45.83 ± 2.14 compared to 50.21 ± 1.75) and increase in NREM sleep time (light-off: 40.16 ± 2.72 vs 33.3 ± 2.64). REM sleep time did not change significantly in NPTX2 KO mice. Our findings demonstrate altered time allocation in Wake and NREM in NPTX2 KO mice.

**Figure 3.**
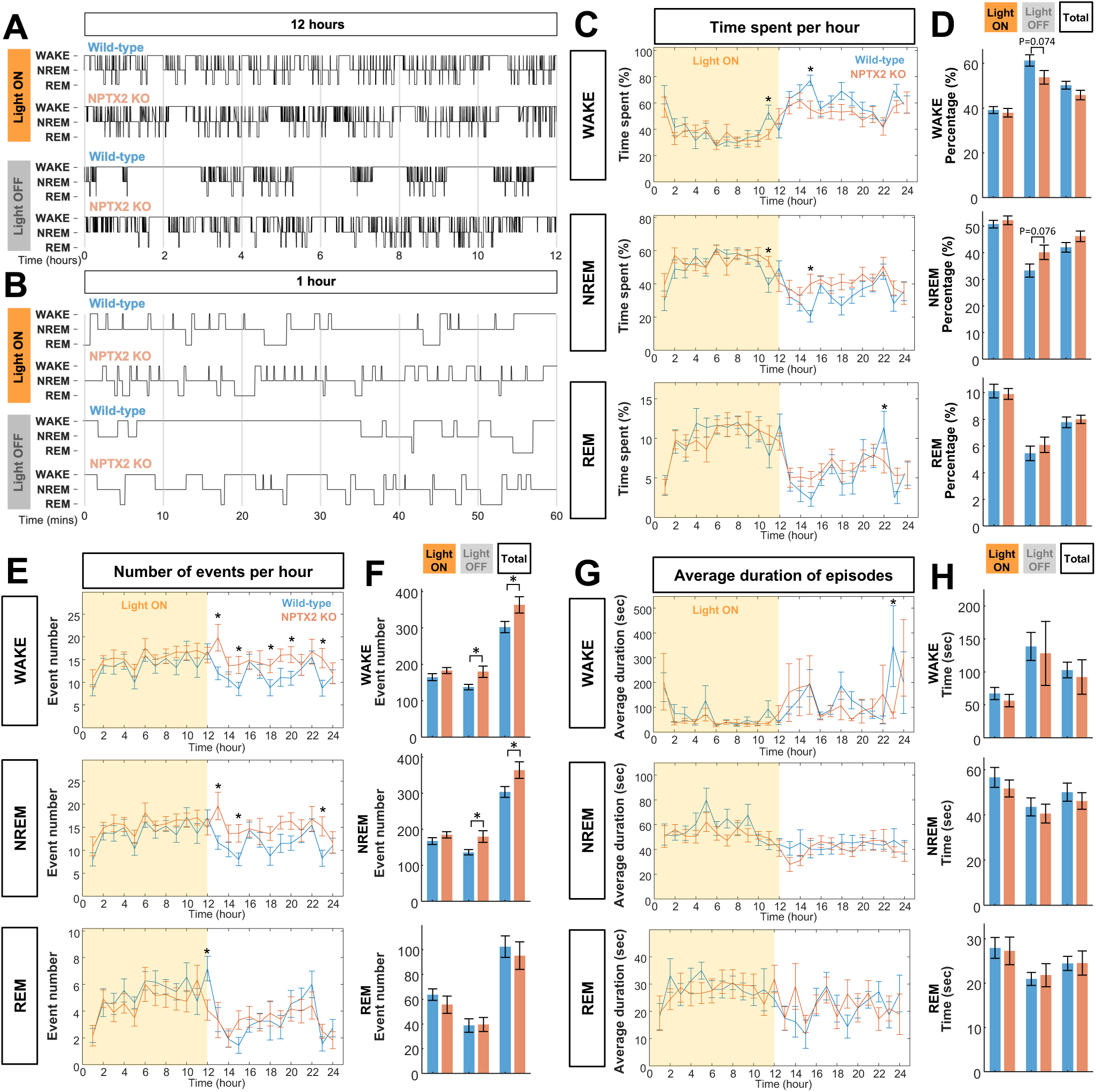
Alterations sleep architecture and sleep fragmentation in NPTX2 KO mice. **(A-B)** Representative hypnograms during 12 hours (A) and 1 hour (B) in wild-type and NPTX2 KO mice for light on and light off phases. **(C)** Hourly percentage of time spent (± SEM) for Wake, NREM and REM across a day of sleep and wake cycle (24 hours) in young wild-type and NPTX2 KO mice. Nonparametric unpaired t-test was performed between the wild-type and NPTX2 KO group for each time point. Significant changes were indicated (*, P<0.05). **(D)** Percentage of time spent (± SEM) for Wake, NREM and REM during light on (12 hours), light off (12 hours) and total (24 hours) comparing wild-type and NPTX2 KO mice. Unpaired t-test was performed between wild-type and NPTX2 KO groups for each phase. P values were indicated. **(E)** Hourly number of events (± SEM) per hour of Wake, NREM and REM across a day of sleep and wake cycle (24 hours) in young wild-type and NPTX2 KO mice. Unpaired t-test was performed between each pair of wild-type and NPTX2 KO group 24 time points. Significant changes were indicated (*, P<0.05). **(F)** Event numbers during light on (12 hours), light off (12 hours) and total (24 hours) comparing wild-type and NPTX2 KO mice. Unpaired t-test was performed between wild-type and NPTX2 KO group for each time range. The significant changes were indicated (*, P<0.05). **(G)** Hourly average duration (± SEM) per hour Wake, NREM and REM across a day of sleep and wake cycle (24 hours) in young wild-type and NPTX2 KO mice. Unpaired t-test was performed between all pairs of wild-type and NPTX2 KO group 24 time points. Significant changes were indicated (*, P<0.05). **(H)** The average duration during light on (12 hours), light off (12 hours) and total (24 hours) comparing wild-type and NPTX2 KO mice. Unpaired t-test was used to compare the wild-type and NPTX2 KO group for each time range. The significant change was indicated (*, P<0.05). N = 11 wild-type and 12 NPT2 KO animals.

To examine sleep fragmentation in NPTX2 KO mice, we conducted further analysis on the number and average duration of episodes for the three vigilance states. Our results demonstrated significant differences in the event numbers per hour for wake, NREM, and REM states between the wild-type and NPTX2 KO mice groups in certain time ranges (Figure 3E). These differences were particularly pronounced around transition time and light-off phase, suggesting that NPTX2 KO mice have difficulties in sustaining wakefulness and or sleep. Additionally, when summarizing the quantification across the sleep and wake cycle, we observed significant differences in the numbers of wake (Figure 3F, light on, P < 0.05; total, P< 0.05) and NREM (Figure 3F, light on, P < 0.05; total, P < 0.05) epochs. Although the changes in hourly average durations of the three vigilance states were not as significant as the number of events, there was a certain time range where it showed a significant increase in Wake (Figure 3G). The representative 12-hour (Figure 3A) and 1-hour (Figure 3B) hypnograms during light-on and light-off phases reflect fragmented sleep and wake bouts in NPTX2 KO mice. These findings indicate a significant level of sleep and wake fragmentation in NPTX2 KO mice, particularly around transition times and light-off phase, suggesting that NPTX2 KO mice experience difficulties with maintaining states and transitioning between day and night, indicating a disrupted sleep architecture.

### Disrupted state transitions and baseline sleep latency in NPTX2 KO mice

Since the circadian rhythm and sleep structure analysis in NPTX2 KO mice indicate that they have difficulties sustaining wakefulness and/or sleep, we investigated whether state transitions are disrupted by analyzing the number of transitions among Wake, NREM, and REM in NPTX2 KO mice across different phases using a custom-written MATLAB script. Similar to the phenotype of altered event numbers and bout duration of Wake and NREM in NPTX2 KO mice (Figure 3), the transitions between Wake and NREM were significantly increased overall. (Figure 4A, light off, P < 0.05 and total, P < 0.05 for both W-N and N-W). These findings further support the notion that NPTX2 KO mice have difficulties in maintaining Wake and NREM states. While transitions from NREM to REM and REM to Wake are common, their reversals are rarely observed. Interestingly, the number of transitions from Wake to REM and REM to NREM was greatly increased in some NPTX2 KO mice showing substantial variation (Supplemental Figure 2), especially in the light-on phase (Figure 4A), although it was not significant in the total NPTX2 KO cohort with an unpaired t-test and a random permutation test (light-on: W-R, P=0.41, R-N, P = 0.66; light off: W-R, P = 0.38, R-N, P = 0.45; total: W-R, P = 0.37, R-N, P = 0.44). However, Wake to REM transitions showed a profound increase in a limited number of NPTX2 KO animals (three out of twelve NPTX2 KOs: 434% increase compared to wild-types, Supplemental Figure 2). This type of transition is one of the major characteristics of narcolepsy, making it an interesting observation. However, it should be noted that our study utilized germ-line NPTX2 KO mice, which might have developed developmental adaptations for the gene deletion. This suggests that the loss of NPTX2 function may enhance these uncommon transitions of Wake to REM and REM to NREM. The analysis of state transitions indicates an increased occurrence of transitions between states, further supporting sleep fragmentation and narcoleptic phenotype in NPTX2 KO mice.

**Figure 4.**
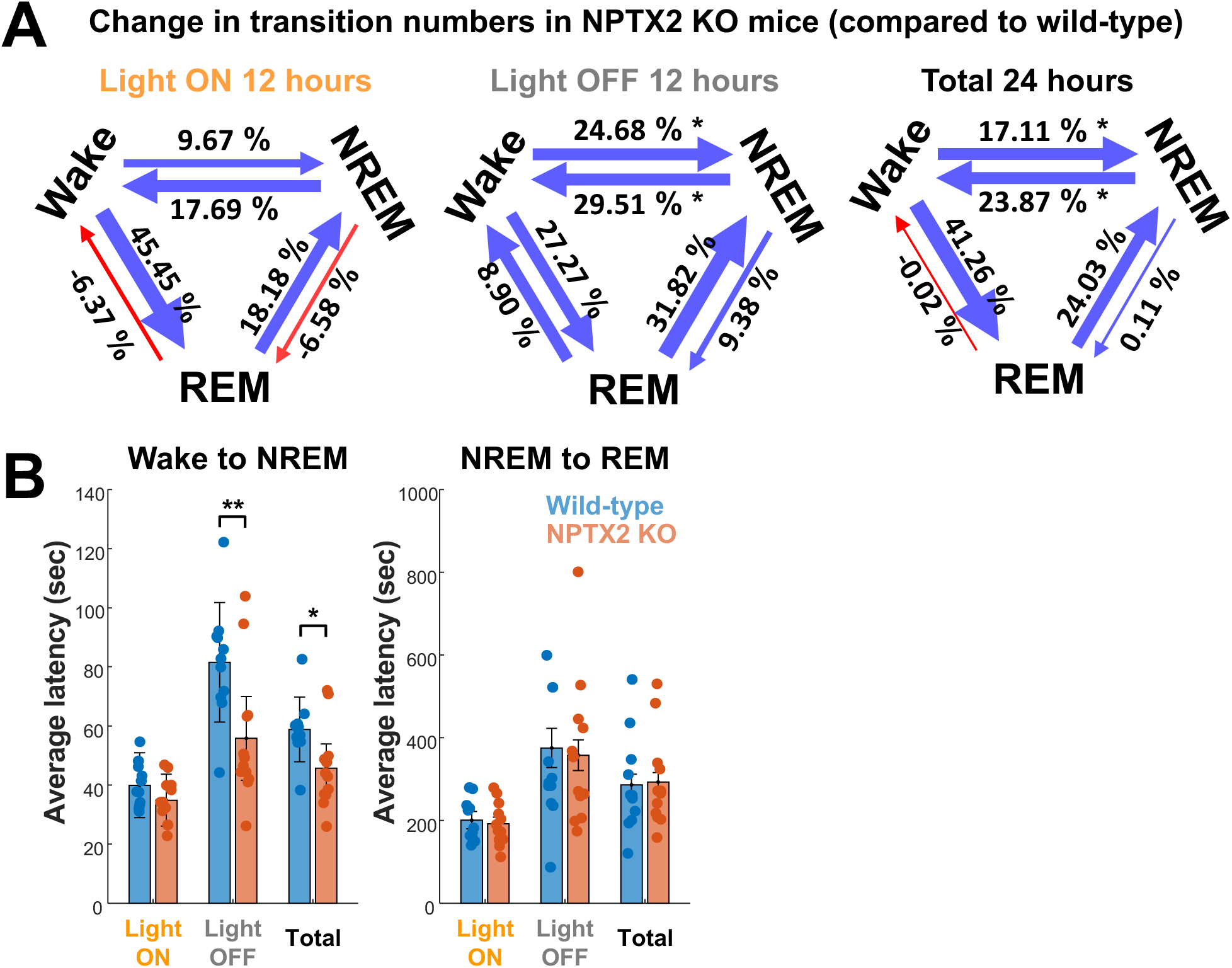
Abnormal state transitions in NPTX2 KO mice. **(A)** Diagrams showing relative changes of transition numbers in NPTX2 KO mice compared to wild-type mice among Wake, NREM and REM in different phases (light on 12 hours, light off 12 hours, and total 24 hours). Blue and red arrows indicate increased and decreased transition numbers in NPTX2 KO mice compared to wild-type, respectively. Changes were indicated as a percentage increase or decrease for each type of transition (Unpaired t-test, *, P<0.05). (B) Sleep latency for NREM and REM comparing wild-type and NPTX2 KO mice during light on, light off and total phases (Unpaired t-test, *, P < 0.05, **, P < 0.01). N = 11 wild-type and 12 NPT2 KO animals.

Consistently, our analysis of baseline latency for Wake to NREM and NREM to REM, time taken to enter the state after the start of the preceding Wake and NREM bouts, reveal that NREM latency is significantly reduced, while not significant during the light-on phase (Figure 4B). NREM to REM latency was not altered in NPTX2 KO mice. The data indicate that NPTX2 KO mice have difficulties in maintaining wakefulness during the light-off phase.

### Altered EEG spectral properties and band power in NPTX2 KO mice

Since clinical sleep disruptions are related to deficits in the broader neuronal network, we analyzed the spectral power of EEG in three distinct vigilance states of NPTX2 KO mice using a custom MATLAB script. To normalize the data, the spectral power of each state was divided by the sum of total power. Our group-level analysis demonstrated significant differences between wild-type and NPTX2 KO mice (Figure 5A, P < 0.0001 for wake, NREM, and REM). Notably, these differences were observed across various frequency bands in all three states, showing an overall reduction of frequency across different bands (Figure 3A). The same analysis during the light-on and light-off phases indicates similar alterations except for enhanced power around theta band in Wake and NREM during the light-off phase compared to the light-on phase (Supplemental Figure 3).

**Figure 5.**
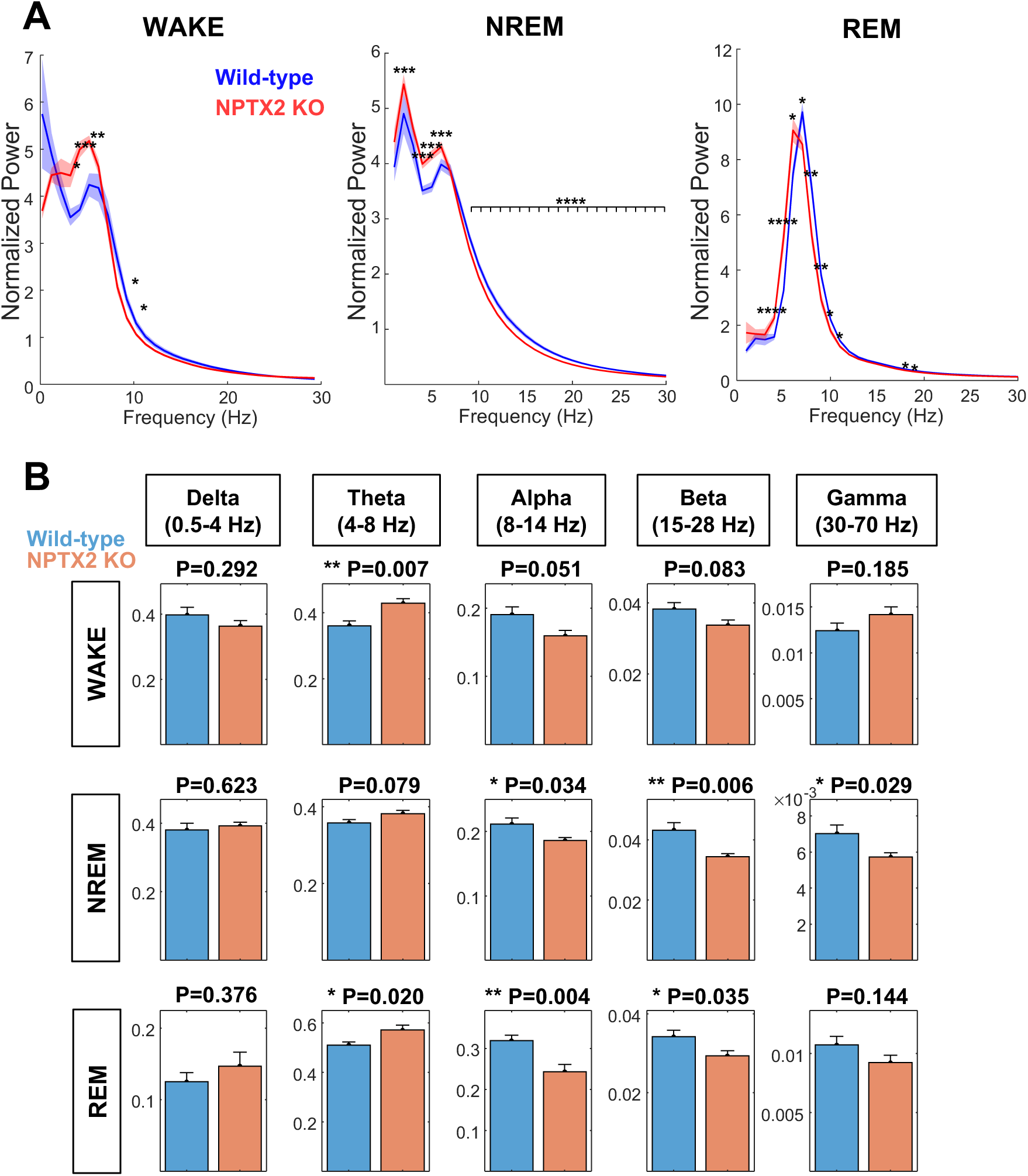
Altered power spectral density and frequency band power during different vigilance states in NPTX2 KO mice. **(A)** Normalized power spectral density (± SEM) from 0 to 30 Hz in 24 hours during WAKE, NREM, and REM, comparing between wild-type and NPTX2 KO mice. Two-way ANOVA was performed for group comparison (P < 0.0001 for Wake, NREM, and REM). Multiple unpaired t-tests were performed to compare the wild-type and NPTX2 KO group for each frequency range. The significant change was indicated (*, P<0.05, **, P<0.01, ***, P<0.001 and ****, P<0.0001). **(B)** Normalized frequency band power (± SEM) of each of the waves (Delta, Theta, Alpha, Beta and Gamma) for WAKE, NREM and REM. Multiple unpaired t-tests were performed to compare the wild-type and NPTX2 KO mice. Significant differences were indicated (*, P<0.05 and **, P<0.01). N = 10 Wild-type and 11 NPT2 KO animals.

Further power analysis focusing on specific frequency ranges, including delta (0.5 - 4 Hz), theta (4 - 8 Hz), alpha (8 - 14 Hz), and gamma (30 - 70 Hz) bands, revealed altered EEG power across multiple bands during wake, NREM, and REM states. Notably, alpha power was mostly significantly reduced in all states, and beta power was decreased during NREM and REM (Figure 5B). The theta power was significantly enhanced in Wake and REM states (Figure 5B). Overall, the shifting tendency is that alpha and beta bands are reduced, and theta is increased. In REM, in particular, theta power exhibited a significant increase, while alpha and beta bands were significantly reduced, indicating substantial shifts to lower frequencies. This observation is intriguing, considering the enriched expression of NPTX2 in Orexin neurons and its potential involvement in stabilizing REM sleep (Feng et al. 2020). Also, interestingly, the theta power was significantly enhanced in NPTX2 KO mice during wake (Figure 5B), and this increase was more pronounced in the light-off phase, but the change was not significant in the light-on phase (Supplemental Figures 3 and 4). Theta oscillation is known to correlate with running speed during spatial navigation (Bender et al. 2015). Hence, enhanced theta oscillation indicates disrupted arousal and locomotion in NPTX2 KO mice that exhibited hyperlocomotive activity upon administration of amphetamine and MK801 (Xiao et al. 2021). These results suggest a systemic deficit in the synchronization of neuronal activity across different vigilance states by NPTX2 loss of function.

### Altered sleep spindle in NPTX2 KO mice

The sleep spindles are strongly related to memory replay, are characteristic oscillatory patterns of EEG activity during NREM, and have high diagnostic and therapeutic implications in neurological diseases such as early AD and schizophrenia (Fernandez and Lüthi 2020). Sleep spindles were shown to negatively correlate with human CSF biomarkers for AD, such as amyloid beta 42, P-tau, and T-tau (Kam et al. 2019). Among them, T-tau showed the most significant association with sleep spindle density. A subsequent study with tau P301S, a mutant tau mouse model, revealed an early alteration in sleep spindle density at 3 months (Kam et al. 2023), whereas the animals display sleep instability at around 10 months (Kam et al. 2023; Holth et al. 2017). Similarly, the reduced level of CSF NPTX2 could predict the progression from MCI to AD (Llano et al. 2023) and even normal cognition to MCI (Soldan et al. 2023), suggesting an early deficit of NPTX2 function. These basic and clinical studies led us to examine the sleep spindle alteration in NPTX2 KO mice. To analyze sleep spindles, we utilized the well-documented algorithm, McSleep spindle detection, which utilizes a mathematical method called sparsity-aware convex optimization (Parekh et al. 2017). Using this method, we could successfully dissociate spindles from transient and residual signals in NREM EEG traces (Figure 6A). The quantification suggests a significant reduction in spindle density and duration in NPTX2 KO mice (Figure 6B). This observation suggests NPTX2 may be critical for memory replay during sleep and may serve as a possible cause for this earliest manifestation in neurodegeneration associated with sleep disturbances.

**Figure 6.**
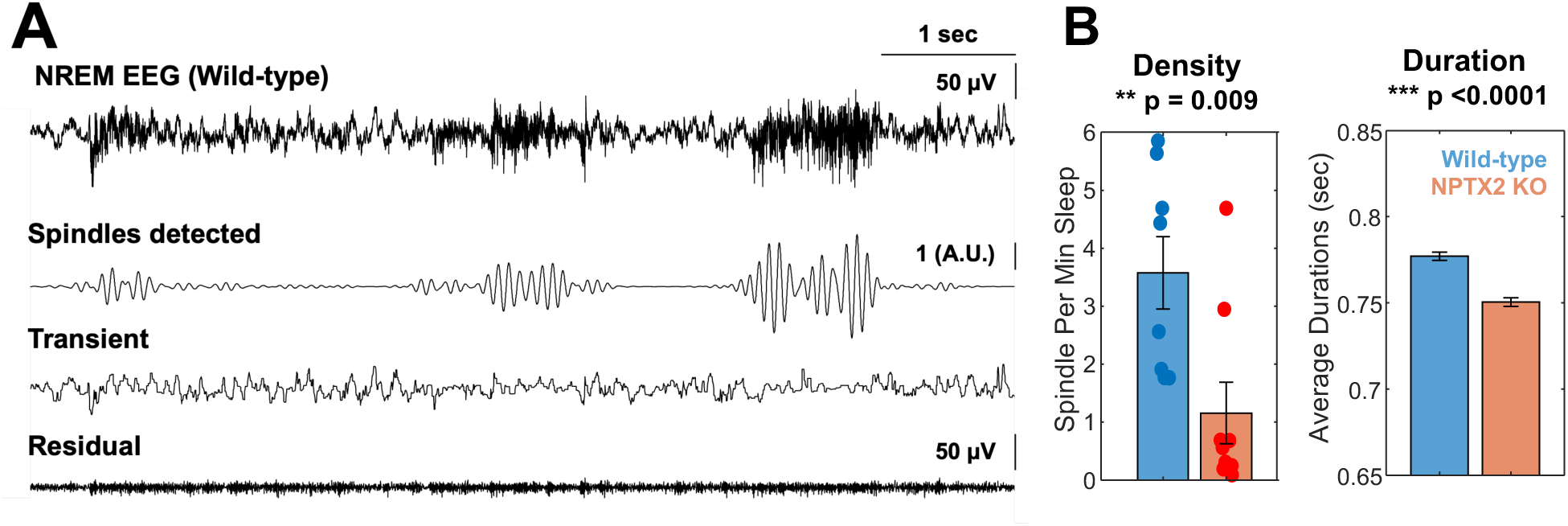
Reduced sleep spindle density and duration in NPTX2 KO mice. **(A)** Representative wild-type NREM EEG trace and spindles detected by McSleep, along with transient and residual signals below. **(B)** Spindle density (spindle number per minute of sleep) and duration of all spindle events (± SEM) were shown. n = 18390 and 12623 events, N= 8 wild-type and 9 NPTX2 KO mice. Unpaired t-test was performed for comparison. P values were indicated.

## Discussion

Based on our observation that reduced NPTX2 is associated with reduced Orexin in the CSF in some normative-aged individuals (Figure 1A) but that Orexin content is retained in NPTX2 KO mice (Figure 1B), we demonstrate that NPTX2 is an effector of sleep and the loss of function contributes to SCRD. The wheel-running behavior study indicates irregular circadian onset time coupled with increased activity during the sleep phase in NPTX2 KO mice, without significant alteration in circadian period (Figure 2). This suggests that they have a relatively normal circadian duration but difficulties in maintaining states, particularly during light transitional periods. Sleep EEG analysis revealed altered time allocation for different vigilance states especially during the light transition period (Figures 3). Sleep and wake fragmentation was evident as revealed by state transition analysis (Figure 4). Moreover, EEG spectral analysis unveils general shifts to lower frequencies of the power spectrum (Figure 5). The systemic changes in EEG power may be attributed to the role of NPTX2 in facilitating the adaptive strengthening of excitatory drive in PV-IN, which subsequently affects neuronal synchrony throughout the brain. Of particular note is the notable enhancement of theta rhythm, which is related to enhanced running speed during spatial navigation (Bender et al. 2015). Additionally, NPTX2 KO mice displayed reduced sleep spindle density, a recognized marker of memory replay and precursor to neurodegenerative change (Figure 6). These findings collectively suggest a potential contribution of NPTX2 loss of function to neurodegenerative and psychiatric diseases, largely mediated by the orchestration of SCRD. Also, they bring into focus the imperative of exploring NPTX2’s involvement in Orexin signaling, serving as a potential conduit between SCRD and the progression of cognitive decline related to aging.

NPTX2 expression is enriched in most Orexin neurons located in the lateral hypothalamus (Reti et al. 2002; Dalal et al. 2013) and NPTX2 is depleted, alongside orexin, in Orexin neurons of human narcolepsy patients (Blouin et al. 2005; Crocker et al. 2005), suggesting a potential interaction between NPTX2 and the Orexin system and contribution to sleep disturbances. The sleep phenotype of sleep-wake fragmentation and alteration in state transitions/sleep latency observed in NPTX2 KO mice shares the sleep disturbance observed in the association between Orexin dysfunction and narcolepsy, a condition characterized by excessive daytime sleepiness (Dauvilliers et al. 2007). Additionally, studies on the Orexin knockout or Orexin degeneration model have demonstrated difficulties maintaining wakefulness and evidence of sleep fragmentation despite relatively normal total amounts of wakefulness (Diniz Behn et al. 2010; Kantor et al. 2009). Also, rats and dogs with impairments in Orexin signaling also exhibit a comparable phenotype, with brief episodes of wakefulness interrupted by short bouts of sleep (Chemelli et al. 1999; Beuckmann et al. 2004). Nonetheless, other prominent indicators of narcolepsy, such as rapid onset of REM, do not appear evident in NPTX2 KO mice (Figure 2B), although they showed increased transitions of Wake to REM in a limited number of animals and altered EEG power in REM (Figure 4A). Also, cataplexy, another feature of narcolepsy, has not been examined in this study. In individuals with narcolepsy, CSF Orexin A and Orexin expression are significantly reduced, with a decrease of approximately 90% (Thannickal et al. 2000; Peyron et al. 2000). Interestingly, however, NPTX2 KO mice displayed increased Orexin contents in the neocortex (Figure 1B), implying that Orexin neuronal degeneration may be upstream of NPTX2 loss in narcolepsy. Hence, it is unclear whether NPTX2 KO mice phenotypes exhibit narcoleptic features. These rather indicate that NPTX2 loss of function is similar to the condition of SCRD during the progression of cognitive decline associated with aging.

NPTX2 is downregulated in individuals with AD, and the CSF levels correlate with cognitive performance and hippocampal volume in MCI/MD (Xiao et al. 2017; Galasko et al. 2019). Reduction of CSF NPTX2 was recently reported to predict the transition from MCI to AD (Llano et al. 2023). Moreover, the reduced level of NPTX2 predicts the progression from normal cognition to the MCI (Soldan et al. 2023). These observations suggest that NPTX2 is one of the earliest measures for detecting cognitive decline during aging. Interestingly, our results indicate that reduced NPTX2 is associated with Orexin reduction in the CSF of aged individuals (Figure 1A), suggesting age-related sleep disturbances are associated with reduced Orexin function which may be modified by NPTX2 reduction. However, the Orexin contents were not reduced in NPTX2 KO mice (Figure 1B), allowing us to examine the specific effect of NPTX2 loss of function on SCRD. Our analysis also revealed that the Orexin in the neocortex was elevated in NPTX2 KO mice which shows the extensive SCRD phenotypes. This implies that further reduction of NPTX2 function during the age-related cognitive decline may contribute to the overproduction of Orexin neurons as compensation for the Orexin neuronal loss, which, in turn, intensifies SCRD phenotypes. Indeed, a recent study suggested that hyperexcitability in Orexin neuronal circuits drives sleep fragmentation during aging in rodent model (Li et al. 2022), which are akin to the SCRD phenotypes of NPTX2 KO mice (Figures 3 and 4). In the neocortex, NPTX2 significantly regulates neuronal excitability by homeostatic modulation of inhibitory interneurons (Chang et al. 2010), and NPTX2 KO primary visual cortex exhibits hyperexcitability (Gu et al. 2013; Pelkey et al. 2015). Further, it was reported that NPTX2 overexpression in Orexin neurons impaired sleep-wake cycle-dependent plasticity of Orexin neuronal axon button number and resulted in melatonin resistance in zebrafish, suggesting that the disruption of NPTX2 rhythmicity can impair structural plasticity of Orexin neurons (Appelbaum et al. 2010). Therefore, it is highly probable that NPTX2 loss of function in the earliest stage of neurodegeneration may drive SCRD by mediating Orexin neuronal dysfunction. This is further supported by reduced sleep spindle density in NPTX2 KO mice (Figure 6), since mishandling of sleep spindle is reported to precede any SCRD symptoms in the neurodegeneration (Kam et al. 2023). Another link between NPTX2 and Orexin may be tau. Interestingly, Orexin is correlated with tau in the CSF of individuals with AD (Liguori et al. 2014). Similarly, the ratio of NPTX2 to tau or p-tau increases the diagnostic performance of the biomarker, suggesting the fundamental relationship between NPTX2 and tau (Galasko et al. 2019). In our analysis, Orexin A is correlated with ptau181 in the CSF of aged individuals (data not shown). These may indicate intricate interplay among NPTX2, Orexin and tau. Therefore, additional research is essential to elucidate the physiological role of NTPX2 within Orexin neurons in the context of SCRD during the aging and cognitive decline. Additionally, a translational study is warranted to directly investigate the connections between sleep disturbances and CSF levels of NPTX2 and Orexin.

How might the deficiency of the IEG NPTX2 contribute to SCRD, potentially implicating Orexin signaling under normal physiological circumstances? The initial stages of NPTX2 deficiency could originate from the cerebral cortex and the hippocampus, areas largely driven by the necessity for maintaining homeostatic equilibrium in response to neuronal activation stemming from wakefulness and experience. Notably, Homer1a has emerged as a central molecular correlate of sleep deprivation, consistently heightening its expression following sleep loss in various mouse models (Maret et al. 2007). Intriguingly, within the same cellular domains where Homer1a is induced, NPTX2 is co-induced, hinting at a potential intertwined function of these genes within the context of the homeostatic response to sleep deprivation. Moreover, recent findings indicate a sequential and hierarchical nature of IEG-mediated memory consolidation, with NPTX2-regulated exocytosis and shedding contingent upon upstream immediate early genes such as Arc and Homer1a (Xiao et al. 2021). Consequently, deficits related to the proper functioning of these genes within the memory consolidation process could potentially lead to NPTX2 deficiency within the cerebral cortex and hippocampus. This, in turn, might trigger augmented excitability in PV-IN, and resultant aberrant activity of postsynaptic PV-IN within the cerebral cortex and hippocampus could disrupt the regulated exocytosis and shedding of NPTX2 within cells, including Orexin neurons, thereby setting the stage for the propagation of NPTX2 deficiency to Orexin neurons. Notably, Orexin neurons form synapses unto PV-IN within the cortex (Aracri et al. 2015; Wenger Combremont et al. 2016). Subsequently, Orexin neurons could experience a reduction in homeostatic inhibition, culminating in a state of hyperactivity and consequently precipitating SCRD. Indeed, Orexin level was elevated in the neocortex of NPTX2 KO mice (Figure 1B). This intricate cascade of events highlights a plausible mechanism by which the deficiency of NPTX2 during aging, orchestrated by the interplay of various IEGs, might trigger a sequence of disruptions that ultimately lead to orexin-mediated SCRD.

## Methods

### Animals

NPTX2 KO mice in congenic C57BL/6J background were obtained from M. Perrin’s laboratory. Animals were kept in ventilated light-controlled boxes where the light is on and off every 12 hours with 24-hour cycle. All procedures were approved and under the guidelines of the Johns Hopkins University Institutional Animal Care and Use Committee.

### NPTX2 and Orexin ELISA assay

#### Human CSF samples

We have used CSF samples of normative-aged individuals from published data set (Alm et al. 2022). CSF NPTX2 was measured by ELISA assay as previously described (Xiao et al. 2017; Xiao et al. 2021). CSF Orexin A was detected using a competitive ELISA Kit (Phoenix Pharmaceuticals) (Liguori et al. 2014).

#### Mouse brain lysate

To determine Orexin levels in NPTX2 KO mice, 2-3 months wild-type and NPTX2 KO mice were euthanized, and dissected cortex, hippocampus and hypothalamus were sonicated in 1% DOC buffer containing 50 mM Tris (pH 9.0), 1% Na-deoxycholate, 50 mM NaF, 20 μM ZnCl_2_, 1 mM Na_3_VO_4_, and protease inhibitor cocktail. Lysates were centrifuged at 16,000 g for 10 min, and the supernatant was used for BCA assay and subsequent Orexin A ELISA assay (Phoenix Pharmaceuticals).

### Electrode implantation surgery or EEG recording

For the EEG recording, we used 2-6 month-old wild-type and NPTX2 KO mice. The EEG electrodes were custom-made in the lab. We soldered tungsten wire (A-M systems) and micro screws which serve as electrodes to an 8-channel pin, which was designed to fit with the Neurologger device. During the surgical procedure, the animals were anesthetized using 4% isoflurane and maintained at a level of 1.2-1.5% throughout the surgery. To prevent swelling and inflammation, a subcutaneous injection of 0.2mg/kg dexamethasone was administered. The skin over the skull was carefully removed, and a recording electrode was implanted into the parietal bone on the right side (AP -5 and ML 1.5 from bregma). A reference/ground electrode was implanted into the occipital bone on the left side (AP -2 and ML -2 from lambda). The pin, electrodes and wires were secured with dental cement (Sun Medical, Japan). Following the surgery, the animals were allowed to recover in their home cages for two days before the recording sessions begin.

### Electroencephalography recording

For the EEG recording, we utilized the Neurologger 2A, a wireless data logger widely used for sleep analysis over multiple days. This device is equipped with its own memory and operates on a battery, eliminating the need for entangled wires that can cause stress to the animals during conventional EEG recording. Out of the 8 channels provided by the Neurologger 2A, two channels were dedicated to serving as the ground and reference, while the remaining channel was used for recording the EEG signal. The EEG signal was recorded at a sampling rate of 1000 Hz with 4X oversampling to ensure high-quality and accuracy of data. In addition to EEG recording, the Neurologger 2A also captured accelerometer data with a high sensitivity of 1000 Hz. For sleep annotation, we utilized this high-density accelerometer data to ensure noninvasive recording for 24 hours (Han et al. 2022). To allow for proper recovery and habituation, the animals were given a two-day recovery period after the electrode implantation and they were exposed to a dummy device that had the same weight as the actual device for 2-3 days. The recording lasted for 24 hours, encompassing both the first 12 hours of light-on period and the subsequent 12 hours of light-off period. To minimize interference, the Neurologger device was attached to the pin on the animals’ head 2-3 hours before the recording began. A timer was set up to ensure that the device started recording at the desired time. After the 24-hour recording period, the data were transferred to a computer using a USB connector for further analysis and processing.

### Sleep annotation and quantitation

Sleep annotation was carried out utilizing custom-written Python scripts that utilized the EEG and accelerometer data collected by the Neurologger device (Lima et al. 2017; Allocca et al. 2019; Han et al. 2022). The scripts implemented a segmentation with 2.5-second intervals, and employed algorithms to approximate the different states based on the characteristics observed in the EEG and accelerometer data. To ensure accuracy and reliability, human annotation was also performed to visually identify and confirm the different brain states, namely wake, NREM sleep, and REM sleep. Our analysis (Figure 1-3) indicates reliable annotation using our system. Using the annotated data, various metrics were computed to analyze sleep efficiency and properties using custom-written MATLAB scripts. Specifically, sleep architecture was analyzed by quantifying time spent, the number of events, and average duration of the three vigilance states in different time bins. Also, sleep transitions among the three states (REM-Wake, REM-NREM, NREM-REM, NREM-Wake, NREM-REM, and Wake-REM), and sleep latency (NREM and RE) were computed and quantified.

### EEG spectral analysis, frequency band analysis and spindle analysis

In order to explore the frequency characteristics of the EEG signals, spectral analysis was conducted using the Fast Fourier Transform (FFT) algorithm using a custom-written MATLAB script. The FFT algorithm was applied to compute the power spectrum of the preprocessed EEG signals. The power spectrum was calculated to obtain the amplitude distribution across different frequency bins (1 Hz bin). To account for individual differences in signal amplitude, the power spectrum was normalized by dividing each frequency bin by the total power within the defined frequency range (1 - 50 Hz). This normalization allowed for a relative comparison of power across different frequency bins.

Using the obtained power spectral density estimates, the frequency range of interest was divided into five distinct bands: delta (0.5 - 4 Hz), theta (4 - 8 Hz), alpha (8 - 14 Hz), beta (15 - 28 Hz), and gamma (30 - 70 Hz). The power within each frequency band was calculated by integrating the power spectral density values within the corresponding frequency range for wake, NREM and REM states and compared between wild-type and NPTX2 KO mice.

Sleep spindles in the 10 – 16 Hz range were detected by a sparse low-rank optimization and duration was determined by McSleep detector that utilizes sparsity-aware convex optimization to separate transients from oscillations (Parekh et al. 2017). We detected sleep spindles from the single-channel EEG recording and quantified the spindle density and duration.

### Wheel running assay and analysis

Wheel-running behavior analysis was performed using ClockLab acquisition software (Actimetrics, IL). 2-4 month old wild-type and NPTX2 KO mice were individually housed in running-wheel cages equipped with sensors that transmitted wheel counts to the acquisition software. Prior to data collection, the mice were acclimated to ventilated light-proof and light-controlled boxes for 2 weeks under a 12-12 hour light-dark (LD) cycle. Subsequently, the animals underwent a recording phase of two weeks under normal LD cycle conditions, followed by two weeks of recording under complete darkness (DD) cycle, and finally, two additional weeks of recording under normal LD cycle conditions. To minimize interference, beddings were carefully changed during the wake period on a weekly basis to avoid inducing movement.

The recorded data, representing the number of wheel revolutions per minute, were analyzed using ClockLab 6 software (Actimetrics, IL). The software’s built-in detection algorithms were utilized to perform analysis of parameters such as period, amplitude, and onset time detection. The average onset delay and onset time coefficient of variation (CV) were calculated from the data collected during the 14-day DD cycle (Figure 3E and F). The onset delay was calculated as the difference in onset time compared to the previous day, and the average onset delay was derived from the 14-day data. The CV, a measure of variability, was calculated by dividing the standard deviation by the sample mean and multiplying by 100. To compare the activity profiles between wild-type and NPTX2 KO mice, the activity data were exported from the software and averaged over the 14 days of LD and DD cycles.

### Statistical analysis

We used Spearman correlation for the analysis of CSF NPTX2 and Orexin A, since the data was not normally distributed, determined by Kolmogorov-Smirnov test. Statistical analyses were conducted to compare the sleep state quantification and spectral analysis data between wild-type and NPTX2 KO mice. Group comparisons were performed using two-way ANOVA to determine significant differences. Posthoc tests, such as unpaired t-tests or Mann-Whitney U tests, were used as indicated in each figure to further analyze specific comparisons. For simple comparisons between wild-type and NPTX2 KO mice, including sleep time, event number and event duration, frequency band analysis, and circadian rhythm parameters, the Mann-Whitney U test was employed. The significance of the results was denoted in each figure with appropriate symbols (*, P < 0.05, **, P < 0.01, ***, P < 0.001, and ****, P < 0.00001). Statistical analyses and data plotting were conducted using either MATLAB or Prism software.

## Supplemental Figure Legends

**Supplemental Figure 1 – related to Figure 1.**
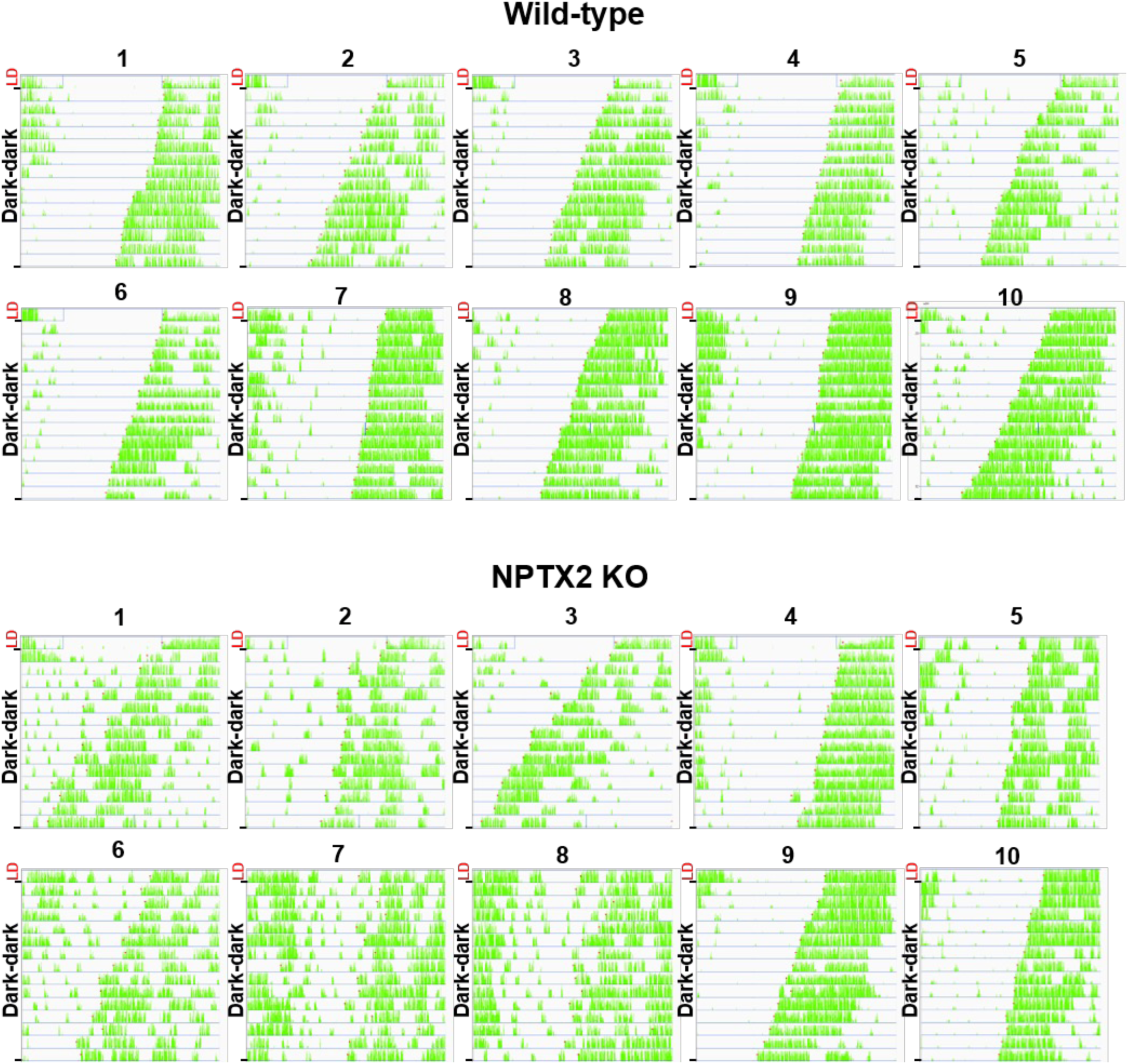
Circadian rhythm onset analysis of all wild-type and NPTX2 KO mice during complete darkness phase. Actograms from one preceding day of the LD cycle and 2 weeks of DD cycle with onset detection (red) marked for each day in 10 wild-type and 10 NPTX2 KO animals.

**Supplemental Figure 2 – related to Figure 3.**
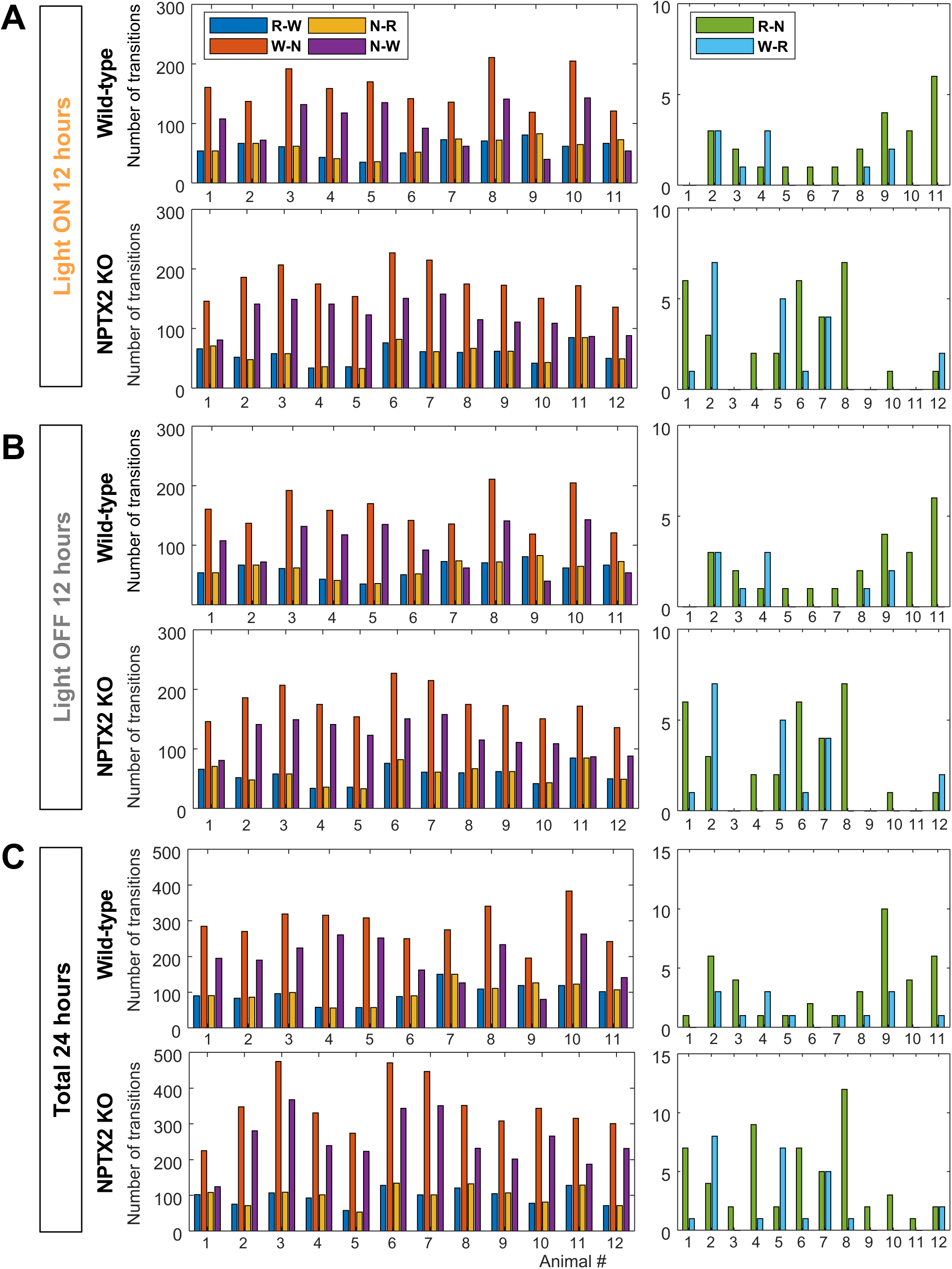
Sleep transitions of all animals. Sleep transition numbers in individual animals for REM-Wake (R-W), NREM-REM (N-R), Wake-NREM (W-N) and NREM-Wake (N-W) (common events, left), REM-NREM (R-N) and Wake-REM (uncommon events, right) in wild-type and NPTX2 KO mice during light on (A), light off (B), and total (C) phases. N = 11 Wild-type and 12 NPT2 KO animals.

**Supplemental Figure 3 – related to Figure 4.**
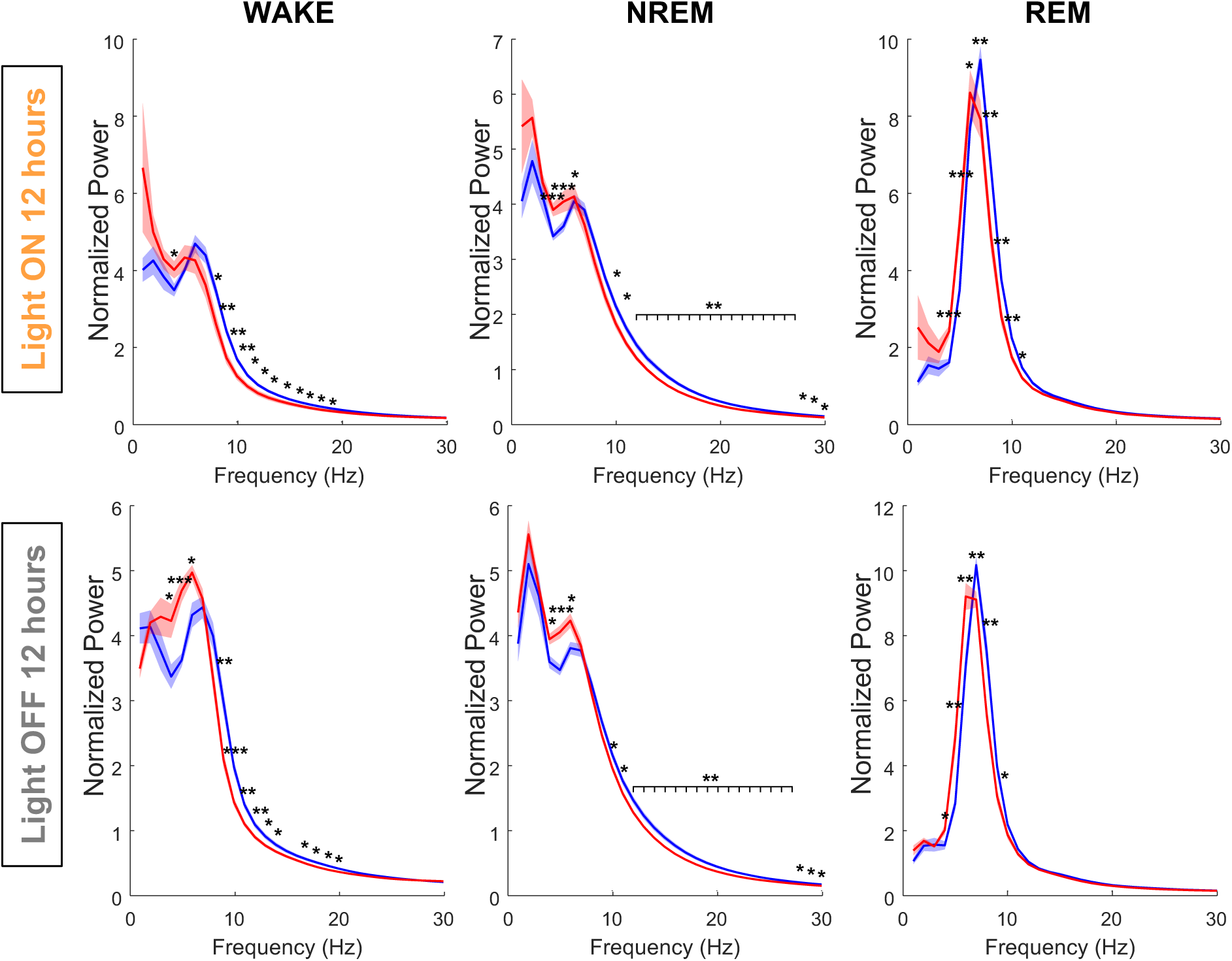
Spectral analysis between light on and light off phases. Normalized power spectral density (± SEM) from 0 to 30 Hz during WAKE, NREM, and REM, comparing wild-type and NPTX2 KO mice between light-on and light-off phases. Two-way ANOVA was performed for group comparison (P < 0.0001 for Wake, NREM, and REM). Multiple unpaired t-tests were performed to compare the wild-type and NPTX2 KO group for each frequency range. The significant changes were indicated (*, P < 0.05, **, P < 0.01, and ***, P < 0.001). N = 10 wild-type and 11 NPT2 KO animals.

**Supplemental Figure 4 – related to Figure 4.**
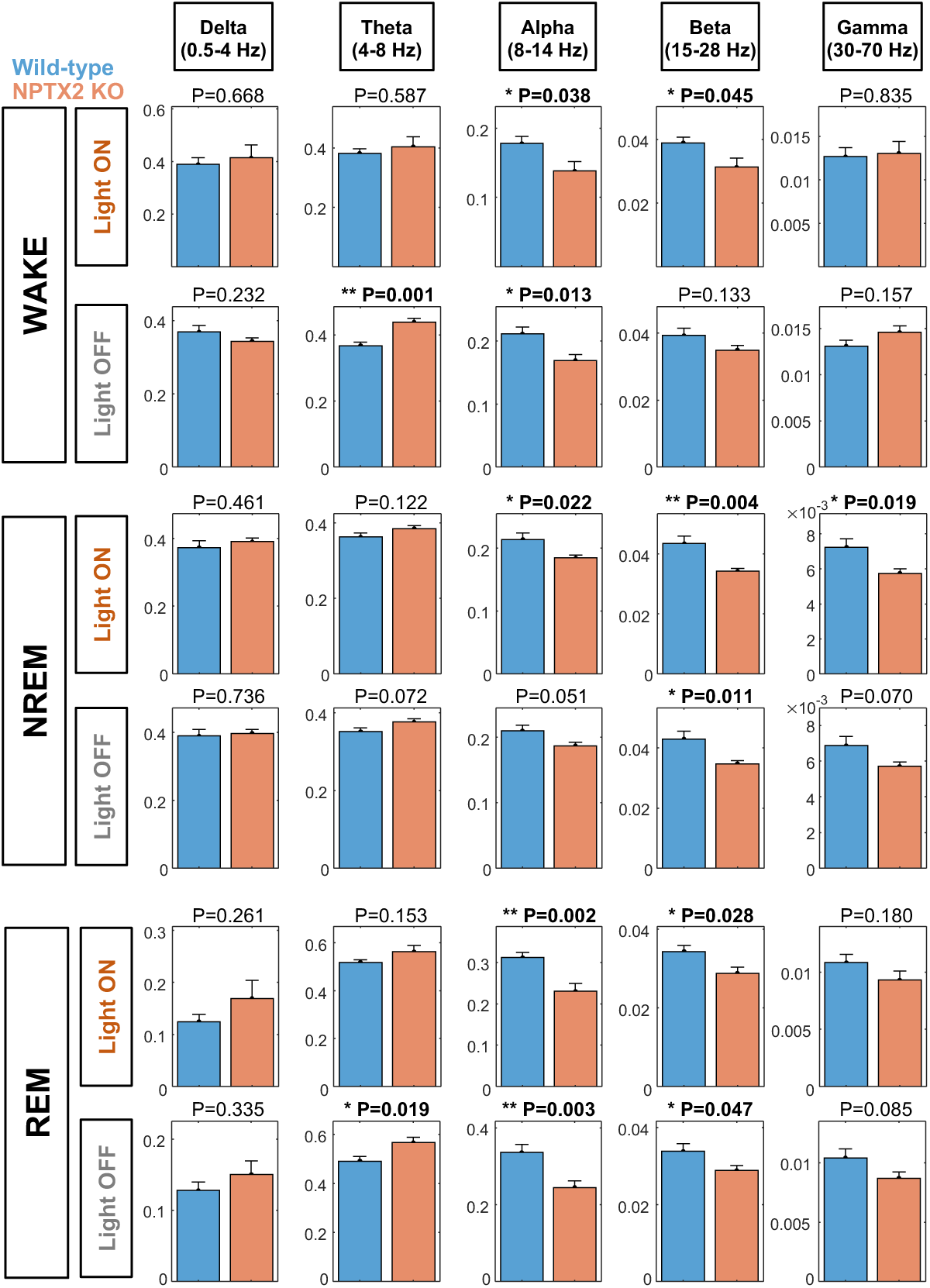
Spectral band frequency analysis between light on and light off phases. Normalized frequency band power (± SEM) of each of the waves (Delta, Theta, Alpha, Beta and Gamma) for WAKE, NREM and REM. Unpaired t-tests were performed to compare wild-type and NPTX2 KO groups per each condition. Significant differences were indicated (*, P<0.05 and **, P<0.01). N = 10 wild-type and 11 NPT2 KO animals.

## Acknowledgments

We thank Dr. Joseph Takahashi, Dr. Mark Wu, Dr. Sangsoo Lee, Dr. Alexei Vitssoski, and Dr. Minhyeok Chang for helpful discussions, Jiali Xiong and Dr. King Yau for advising ClockLab analysis. This work was supported by R35NS097966 NIA/NINDS to P.W., P01AG009973 NIH/NIA to P.W. and P30AG066507 NIH/NIA to Johns Hopkins University (JHADRC Junior Faculty Award).

## Author Contributions

The study was conceived and designed by S.R. and P.W. EEG recording and wheel-running experiments were conducted by S.R. and A.D. S.R. performed the data analysis. C.K. developed the Python script for semi-automatic sleep annotation and quantification. S.R. wrote MATLAB scripts for sleep quantitation, state transition analysis, EEG spectral analysis, and spindle analysis. M.X. performed ELISA. A.S. assisted with the Neurologger EEG recording. H.K. provided analytic resources. A.B. provided human CSF samples and collected information. The draft manuscript was written by S.R. and finalized by S.R. and P.W.

